# Disentangling environmental drivers of circadian metabolism in desert-adapted mice

**DOI:** 10.1101/2020.12.18.423523

**Authors:** Jocelyn P. Colella, Danielle M. Blumstein, Matthew D. MacManes

## Abstract

Metabolism is a complex phenotype shaped by natural environmental rhythms, as well as behavioral, morphological, and physiological adaptations. Although historically studied under constant environmental conditions, continuous metabolic phenotyping through environmental transitions now offers a window into the physiological responses of organisms to changing environments. Here, we use flow-through respirometry to compare metabolic responses of the desert-adapted cactus mouse (*Peromyscus eremicus*) between diurnally variable and constant environmental conditions. We contrast metabolic responses to circadian cycles in photoperiod, temperature, and humidity, against those recorded under constant hot-and-dry and constant cold-and-wet conditions. We found significant sexual dimorphism in metabolic responses, despite no measurable difference in body weight. Males seem to be more heat tolerant and females more cold tolerant. Under circadian environmental cycling, the ratio of CO_2_ produced to O_2_ consumed (the respiratory quotient or respiratory exchange ratio) reached greater than one, a pattern that strongly suggests that lipogenesis is contributing to the production of energy and endogenous water in this species. This hypothesis is consistent with the results of previous dehydration experiments in this species, which documented significant weight loss in response to dehydration, without other physiological impairment. Our results are also consistent with historical descriptions of circadian torpor in this species (torpid by day, active by night), but reject the hypothesis that this pattern is driven by food restriction or negative water balance, as both resources were available to animals throughout the experiments.

**SUMMARY STATEMENT:** Continuous metabolic phenotyping of desert-adapted cactus mice (*Peromyscus eremicus*) identifies significant metabolic differences between the sexes and circadian patterning consistent with lipogenesis and environmental entrainment.

## INTRODUCTION

Our planet is undergoing unprecedented environmental change, which is exerting new, strong selective pressures on species, forcing them to relocate, adapt *in situ,* or risk extinction. Warming temperatures and increased desertification are predicted to impact most species in North America (Parmesan and Yohe, 2003; Urban, 2015; IPCC, 2018). One productive line of research has been related to desert animals (Greenville et al., 2017; Kordonowy et al., 2017; MacManes, 2017; Riddell et al., 2019) which are already adapted to hot and dry conditions and therefore may illustrate mechanisms (*e.g.,* molecular, behavioral, or physiological) through which animals may adapt to a warming climate (Vale and Brito, 2015; Colella et al., *in press*). Much research has been devoted to understanding genetic correlates with heat- and dehydration-tolerance in extant desert-adapted taxa. Of equal importance, are the physiological adaptations of desert species, which are predicted to play an important role in species responses to climate change (Kearney et al., 2009; Boyles et al., 2011; Huey et al., 2012). Yet, adaptive thermoregulatory mechanisms do not evolve under fixed (or static) environmental conditions, as they have traditionally been measured (Hulbert and Else, 2004). Instead, thermoregulatory responses are also shaped by natural environmental rhythms, including circadian and seasonal variation in temperature, humidity, and photoperiod, among other variables (Morrison et al., 2008; Seebacher, 2009). Circadian rhythms are observed in most mammals and are in part governed by genetics, as well as the interplay between the perception of environmental change and neuroendocrine-mediated physiological responses (Miranda-Anaya et al. 2019). Examination of the physiological responses of organisms to both static and dynamic environmental conditions provides a window into the flexibility of thermoregulatory responses to changing environments and will be increasingly important to understanding, and ultimately, predicting (Garland and Carter, 1994; Dalziel et al., 2009; Easterling et al., 2000; IPCC, 2012; Huey et al., 2012) species responses to climate change.

Deer mice of the genus *Peromyscus* inhabit an exceptional range of thermal environments in North America, ranging from cold high elevation mountain tops to arid hot deserts (Hock, 1964; Cheviron et al., 2014; MacManes and Eisen, 2014; Harris and Munshi-South, 2017; Storz et al., 2019). Under projected climate scenarios (e.g., IPCC, 2018), adaptations that increase fitness in warmer, drier environments are predicted to be increasingly favored (Merkt and Taylor, 1994; Gienapp et al., 2008; Kingsolver, 2009). Cactus mice *(Peromyscus eremicus)* are endemic to the southwestern deserts of North America and exhibit a number of morphological (large ears, large surface to volume ratio), behavioral (saliva spreading, Murie, 1961), and physiological (low metabolism, Murie, 1961, McNab and Morrison, 1963; anuria, Kordonowy et al., 2017) adaptations to high heat, low water desert environments. The suitability of this species to life in captivity also makes them a promising experimental model for identifying and characterizing thermoregulatory mechanisms.

In addition to being able to withstand environmental extremes, desert taxa, including cactus mice, are also adapted to large circadian fluctuations in environmental conditions. In the Sonoran Desert, daily temperatures can swing between −4°C to 38°C (Reid, 1987; Sheppard et al., 2002), with weeks or even months passing between rainfall events. Water is required for basic physiological functioning (Haussinger, 1996; Montain et al., 1999), but in desert environments water loss often exceeds intake (Heimeier et al., 2002) and evaporative water loss is further exacerbated by extreme heat and aridity (Cheuvront et al., 2010). Most endotherms are limited in their ability to metabolically respond to sub-optimal environmental conditions (Bennet and Ruben, 1979; Fuglei and Oritsland, 1999), with a notable exception being those that can enter hibernation or temporary torpor (Ruf and Geiser, 2015). In response to water stress, cactus mice enter seasonal torpor, reducing above ground activity and depressing metabolism during the dry season (Macmillen, 1965). Early investigations found that torpor in cactus mice could be initiated at any time by food restriction, but could only be triggered by dehydration during the summer months, which suggests some degree of metabolic entrainment (Macmillen, 1965, 1972). As a nocturnal species, cactus mice also reduce activity levels during the daytime, seeking cooler, shaded parts of their habitat, hiding underground, or even entering circadian torpor during the day (torpid by day, active by night) to decrease heat load, respond to resource limitation, maximize heat dissipation, and minimize respiratory water loss (Macmillen, 1965, 1972; Degen, 2012; Alonso et al., 2016). Ultimately, a balance exists between environmental heterogeneity *(e.g.,* food availability, thermal stress), which requires animals to sense and respond to external conditions in real time via reactionary mechanisms, and environmental predictability which can lead to the evolution of anticipatory biorhythms based on regular abiotic conditions *(e.g.,* photoperiod).

Here, we examine patterns of metabolic variation in cactus mice across a 24-hour period in response to both diurnally variable and constant environments. We hypothesize that cactus mice will exhibit physiological responses consistent with desert adaptation, that minimize daytime energy expenditure and water loss. We further hypothesize that cactus mouse metabolism will be well-tuned to circadian patterns of environmental variation, and that circadian patterning will be disrupted under constant environmental treatments. Comparing metabolic responses between sexes and across constant and environmentally variable treatment groups, we show that (i) males are more heat tolerant than females, (ii) patterns of metabolic variation are well-tuned to environmental cycling, and (iii) lipogenesis offers a potential mechanism of endogenous water production, only employed under variable environmental conditions.

## MATERIALS AND METHODS

### Mice & environmental conditions

Mice were cared for, handled, and sampled in accordance with the University of New Hampshire’s Institutional Animal Care and Use Committee (AUP #180705). All mice used in this study were bred from wild-derived lines maintained at the *Peromyscus* Genetic Stock Center of the University of South Carolina (Columbia, South Carolina, USA). All mice were sexually mature, non-reproductive adults between three and nine months of age. Animals were fed LabDiet 5015* (19% protein, 26% fat, 54% carbohydrates, food quotient [FQ] of 0.89) and provided water *ad libitum* from a well-sealed water bottled. Body weight (g) was collected for each animal at the start of each experiment. All mice were non-reproductive adults and were housed alone, with bedding, and acclimated to experimental cages and conditions for 24-hours prior to metabolic measurement. Environmental conditions for each treatment illustrated in Fig. 1. In brief, the desert simulation chamber has a photoperiodic cycle of 16 hr of light and 8 hrs of darkness, with photoperiod transitions occurring at 06:00 (dark to light) and 20:00 (light to dark) to mimic sunrise and sunset. Photoperiod remained constant across all treatments. To mirror normal (baseline) circadian variation of Southwestern deserts, the environmental chamber was held at a daytime temperature of 32°C and relative humidity (RH) of 10% from 09:00 to 20:00. Chamber temperature and humidity were continuously recorded through the Sable Systems International (SSI; Las Vegas, NV, USA) Field Metabolic System (FMS). At 20:00, the chamber temperature is decreased and humidity increased gradually over the course of one hour to 24°C and 25% RH, respectively. This one hour transition period (20:00-21:00) is hereafter referred to as the evening transition (T2), whereby the room shifts from hot-and-dry to cool-and-wet following lights-out at 20:00. After 21:00, conditions were held constant overnight until 06:00. This period is hereafter referred to as nighttime. At 06:00, following sunrise (e.g., lights on), there is a gradual 3-hour transition from cool-and-wet nighttime conditions (24°C and 25% RH) to hot-and-dry daytime conditions (32°C and 10% RH), during what is hereafter referred to as the morning transition period (T1). We tested two constant temperature treatments: the hot treatment maintained daytime temperature and humidity (32°C and RH of 10%) levels throughout the entire experiment and the constant cold treatment maintained nighttime environmental conditions (24°C and 25% RH).

**Figure 1.**
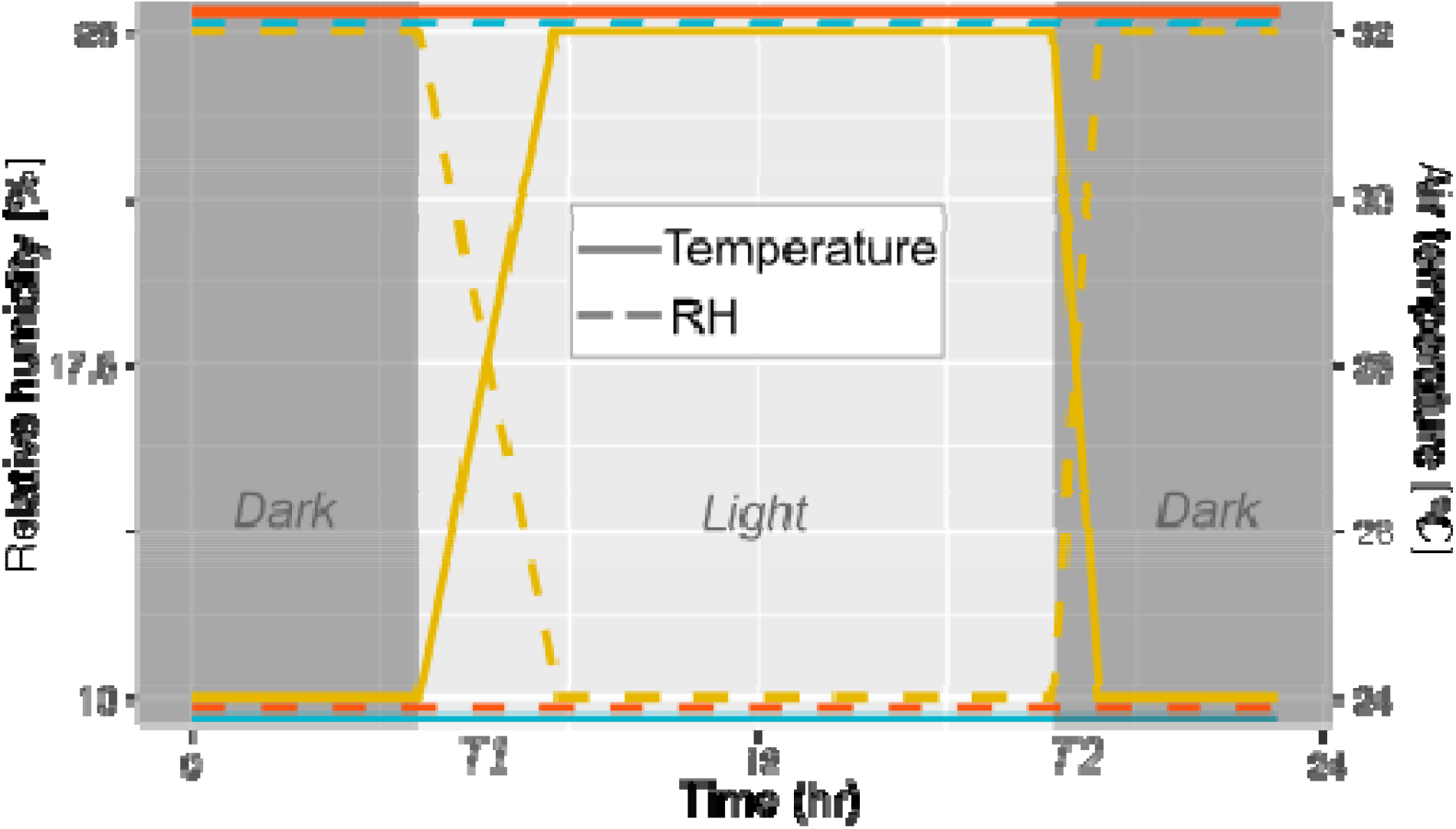
Schematic of environmental variables across a 24-hour cycle for each treatment group: diurnally variable baseline treatment in yellow, constant hot treatment in orange, and constant cold treatment in blue. Temperature is indicated by a solid line and relative humidity (% RH) by dashed lines. Photoperiod is indicated by light and dark grey boxes.

### Metabolic phenotyping

Metabolic variables were measured using an 8-cage FMS manufactured by SSI. We recorded oxygen consumption (VO_2_), carbon dioxide production (VCO_2_), and water vapor pressure (WVP, kPa) over 72-hours for an empty baseline chamber (e.g., ambient air) and seven animal chambers with an equivalent volume of 9.5 L. Air was pulled from chambers at a mass flow rate of 1600 ml/min (96 L/hr), multiplexed through the SSI MUXSCAN multiplexer, and sub-sampled at 250 ml/min, per the SSI FMS manual. A flow rate of 1600 ml/min and chamber volume of 9.5 L equate to a time constant of 5.9 minutes to sample and replace the entire container volume (Lighton, 2017). Chamber sampling alternated between a baseline chamber measurement for 120 s followed by a random animal chamber measurement for 120 s each, resulting in approximately two measurements per animal per hour. Frequent baseline measurements were important for reducing measurement noise introduced by environmental transitions. This protocol was repeated twice for each sex under each treatment (baseline/diurnal, hot, cold), resulting in three days of continuous metabolic measurements for 14 adult males and 14 adult females for each treatment. Barometric pressure (BP, Torr), temperature (degrees Celsius [°C]), and relative humidity (RH, %) of the environmental chamber were recorded throughout. Raw data were processed using two macros (available on Dryad) in the ExpeData analytical software (SSI). The first macro step calculated VO_2_, VCO_2_, and VH_2_O (ml min^-1^ at standard temperature and pressure [STP]) with smoothing, z-transformation, lag correction (O_2_/CO_2_ = 18 s lag, H_2_O = 12 s lag), and gas measurements and flow rates adjusted for BP and WVP (humidity). The second macro step averaged chamber measurements across the most stable 50% of each 120 s measurement window to produce a single average value for each metric within each window. VO_2_ was used as a proxy for metabolic rate (MR), per Hulbert and Else (2004). As a measure of metabolic fuel use, the respiratory quotient (RQ, L/hr), also called the respiratory exchange rate (RER), was calculated as the ratio of VCO_2_ to VO_2_ (Lighton, 2017). Energy expenditure (EE, kcal/hr), or the sum of the basal metabolic rate (BMR), physical activity (typically, 4-16%; Abreu-Vieira et al., 2015), and cold-induced thermogenesis, was calculated as in Lighton (2017, eq. 9.15). BMR is difficult to measure in mice due to the confounding effects of activity, constant flux in postabsorptive state, and variations in body temperature, which were not directly measured in this study (Abreu-Vieira et al., 2015). Measurements located more than three standard deviations away from the mean for each response variable were identified as outliers, likely non-biological in nature, and were removed from downstream analyses.

### Statistical Analysis

All analyses were conducted in *R* v 3.6.1 (R Core Team, 2012), unless otherwise specified. All code is freely available at: www.github.com/jpcolella/peer_respo. After first confirming that the data were normally distributed *(shapiro_test* function in the package *rstatix* v. 0.6.0, Kassambara, 2020) and had homogeneous variance *(stats::var.test),* we ran an unpaired two-sample t-test *(stats::t.test,* p < 0.05) to test for sexual dimorphism in weight. Data were subset by sex, experiment, and time interval (morning transition, daytime, evening transition, nighttime) to generate summary statistics. We calculated the mean (+/− sd), median, and range (minimum, maximum) for each time interval, experiment, and sex. We used a student’s two-tailed t-test (*t.test*), adjusted for multiple comparisons (Bonferroni correction, *p.adjust*), to test for significant (p < 0.05) differences in response variables between the sexes under each experimental condition. Significant differences in male and female responses (Fig. S1 and S2) led us to analyze each sex independently in all downstream analyses.

To determine if weight was a covariate, we ran a general linear regression between weight and each response variable to test for a significant relationship (R^2^, p < 0.05). We found that weight was not a significant predictor of metabolic response and thus did not include it as a covariate (Fig. S3). We evaluated ANOVA assumptions in R, including: independent observations, no significant outliers, data normality, and homogeneity of variance. Due to ANOVA assumption violations, we elected to run a Kruskal-Wallis test (*stats::kruskal.test*) as a non-parametric alternative to a one-way ANOVA (Hollander and Wolfe, 1973). The Kruskal-Wallis test extends the two-sample Wilcoxon test to situations where there are more than two groups, to identify significant differences in group means across experimental treatments (p < 0.05, Bonferroni corrected).

### Time-series analyses

Time-series data were visualized in *ggplot2* v. 3.3.0 (Wickham, 2016) and *visreg* v. 2.7.0 (Breheny and Burchett, 2017) across a 24-hour period using both local regressions, as a non-parametric approach that fits multiple regressions in a local neighborhood *(loess* option in *ggplot2),* and a generalized additive model (GAM; Wood, 2011) in the *mgcv* package v. 1.8-31 (Wood 2004, 2017; formula: response variable ~ s(time), where time is measured in seconds across a 24-hour cycle). Plots were generated in R and edited in the free, open-source vector graphics editor InkScape (https://inkscape.org). Distributions were contrasted across treatment groups.

Change point analyses were used detect significant transitions in the mean and variance *(changepoint::cpt.meanvar*) of each response variable across a 24-hour cycle *(changepoint* package v. 2.2.2, Killick and Eckley, 2014; Killick et al., 2016). Change points identified under baseline conditions were compared to those detected under constant hot and cold treatment groups to determine whether organismal responses were anticipatory or reactive relative to environmental change and whether these shifts remained unchanged or not across experimental conditions. Because additional parameters almost always improve change point model fit, it is important to quantitatively identify meaningful penalty thresholds to avoid overparameterization (Killick and Eckley, 2014; Haynes et al., 2017). Selection of penalty thresholds depends on many factors, including the size and duration of change, which remain unknown prior to change point analysis (Guyon and Yao, 1999; Lavielle, 2005; Birge and Massart, 2007). Current best practices are based on visual inspection of change point estimates compared to time series data and known forcing functions, if available (Killick and Eckley, 2014). For example, there are two known (artificially controlled) environmental transitions under the baseline or diurnally variable treatment conditions; therefore, a minimum of two change points, maximum of four (corresponding to the beginning and end of each transition), are expected and additional or fewer identified change points should be interpreted with caution and explored in greater detail (Killick and Eckley, 2014). Sharp transitions in slope for example are especially difficult to fit into discrete bins, and may cause additional change points to be estimated within steep environmental transitions that may not be biologically relevant (Fearnhead et al., 2019). To estimate penalty thresholds, we used the CROPS (Change points for a Range Of Penalties; Haynes et al., 2017) penalty setting with the PELT (Pruned Exact Linear Time; Killick et al., 2012) method to test a range of penalty values (λ, 5-500) and to optimize computational efficiency. To balance improved model fit with the addition of more parameters, we plotted the difference in the test statistic versus the number of change points detected to identify an optimal penalty threshold of 20 as appropriate for all but one comparison (e.g., Δ log-likelihood > λ). Diagnostic plots for baseline male water loss did not asymptote with the x-axis under a penalty threshold of 20, thus we used a penalty value of 18 for this test. Under the assumption that a large improvement in model fit is indicative of a true change point (Lavielle, 2005; Killick, 2017), we used Akaike information criterion (AIC) diagnostic plots to identify the optimal number of change points based on our data (maximum-likelihood). Visual inspection of changepoints plotted over our raw data allowed us to identify change points that may be over fit. As a Bayesian alternative, we also used the *bcp* package v. 4.0.3 (Erdman and Emerson, 2007) to estimate change points based on a posterior probability threshold of 0.2. The occurrence time of significant change points were compared across treatment groups and against known room transitions to estimate lag-times and identify anticipatory versus reactionary metabolic responses. The mean and slope of each segment was calculated in R.

## RESULTS

### Metabolic phenotyping

In total, we recorded 13,020 120-s intervals (6,550 female, 6,470 male). We measured 4,020 baseline observations (2,038 female and 1,982 male). We measured metabolic variables for 4,506 distinct 120-s intervals under the constant cold treatment: 2,256 female and 2,250 male measurements. For the constant hot treatment, we measured 4,494 total observations (2,256 female, 2,238 male). After removing outliers, we retained 6,335 female measurements total: 1,877 measurements under baseline conditions, 2,230 under constant hot conditions, and 2,228 under constant cold conditions. We retained 6,255 measurements for males: 1,844 baseline, 2,214 hot, and 2,197 cold. Female weights ranged from 15.4 to 25.2 g and male weights ranged from 15.5 to 26.6 g. All raw data (Expedata files) and processed machine-readable csv files are available on Dryad: https://doi.org/10.5061/dryad.f4qrfj6v0.

### Static statistics

We found no significant difference between mean weights for males and females, and no relationship between weight and any measured metabolic variable (Fig. S3). T-tests between the sexes identified significant differences in metabolic responses, with few exceptions (Table 1; Fig. 2; Figs. S1 and S2). Summary statistics for all metabolic response variable are reported in Table 1 for each sex and experimental group (baseline, hot, cold). Mean EE was higher for males under all conditions, while mean water loss was only higher for males under the cold treatment. Under diurnal conditions, RQ patterning was similar among the sexes with both males and females reaching an RQ greater than 1 around midday and RQ was less than the FQ overnight (Table 1). Under constant cold conditions both sexes exhibit a fixed RQ just above the FQ, while under constant hot conditions female RQ was fixed and equal to the FQ, while male RQ was slightly above the FQ (0.94; Table 1) but also fixed.

**Figure 2.**
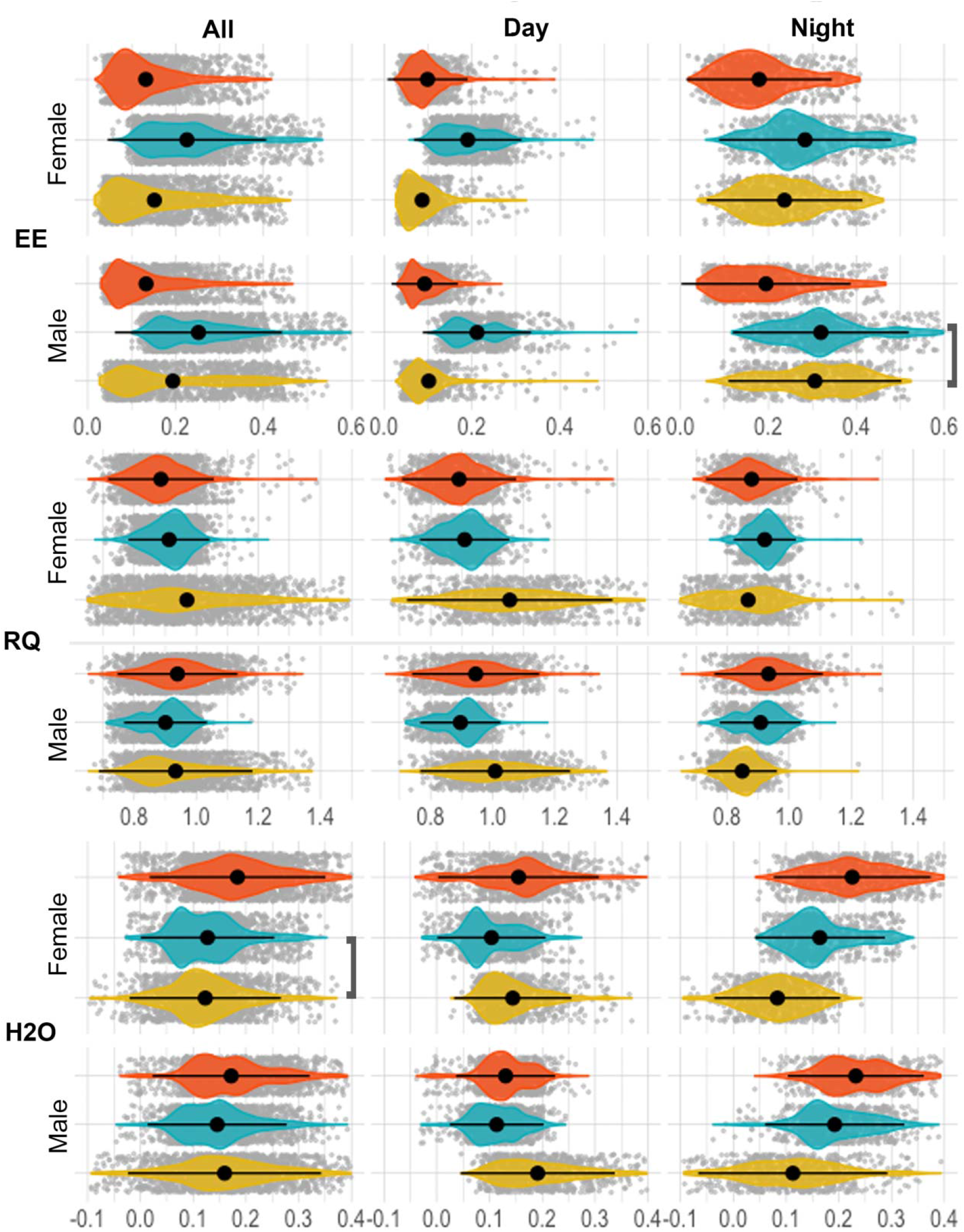
Violin plots showing mean (black dot) for energy expenditure (EE), respiratory quotient (RQ), and water loss (H_2_O). Standard deviation is shown as a horizontal black line and all observations are represented by grey dots. All but two group means are significantly different (Bonferroni corrected, Wilcoxon test, p < 0.05). Left column: entire 24-hour cycle (All); Central: daytime only; Right: nighttime only. Gold = diurnally variable baseline, blue = cold, orange = hot. The only two insignificant comparisons are labeled with a bracket.

**Table 1.** Summary statistics (median = med., mean, standard deviation = s.d., range) for females (top) and males (bottom) for each response variable: energy expenditure (EE), water loss (H_2_O, mg), respiratory quotient (RQ), carbon-dioxide produced (VCO_2_), and oxygen consumed (VO_2_) across each experiment (Baseline, Cold, Hot) during the daytime (rest) and nighttime (active) periods.

|  | Data | EE |  |  |  | H2O (mg) |  |  |  | RQ |  |  |  |
| --- | --- | --- | --- | --- | --- | --- | --- | --- | --- | --- | --- | --- | --- |
|  |  | med. | mean | sd | range | med. | mean | sd | range | med. | mean | sd | range |
| Females | All | 0.15 | 0.17 | 0.10 | 0.02-0.53 | 0.14 | 0.15 | 0.08 | -0.11-0.42 | 0.91 | 0.92 | 0.12 | 0.51-1.50 |
|  | Baseline | 0.12 | 0.15 | 0.10 | 0.02-0.46 | 0.12 | 0.12 | 0.07 | -0.11-0.37 | 0.94 | 0.97 | 0.18 | 0.51-1.50 |
|  | Day | 0.08 | 0.09 | 0.05 | 0.03-0.32 | 0.13 | 0.14 | 0.06 | 0.03-0.37 | 1.05 | 1.05 | 0.17 | 0.62-1.49 |
|  | Night | 0.23 | 0.24 | 0.09 | 0.04-0.46 | 0.09 | 0.08 | 0.06 | -0.11-0.24 | 0.87 | 0.86 | 0.12 | 0.52-1.36 |
|  | Cold | 0.21 | 0.23 | 0.09 | 0.06-0.53 | 0.12 | 0.13 | 0.06 | -0.03-0.35 | 0.92 | 0.91 | 0.07 | 0.67-1.23 |
|  | Day | 0.18 | 0.19 | 0.06 | 0.06-0.48 | 0.09 | 0.10 | 0.05 | -0.03-0.27 | 0.92 | 0.91 | 0.07 | 0.67-1.18 |
|  | Night | 0.27 | 0.28 | 0.10 | 0.06-0.53 | 0.15 | 0.16 | 0.06 | 0.04-0.34 | 0.93 | 0.92 | 0.05 | 0.74-1.23 |
|  | Hot | 0.11 | 0.13 | 0.07 | 0.02-0.42 | 0.18 | 0.19 | 0.08 | -0.04-0.42 | 0.88 | 0.88 | 0.09 | 0.51-1.39 |
|  | Day | 0.09 | 0.10 | 0.05 | 0.02-0.39 | 0.16 | 0.16 | 0.08 | -0.04-0.41 | 0.89 | 0.89 | 0.10 | 0.51-1.39 |
|  | Night | 0.17 | 0.18 | 0.08 | 0.02-0.41 | 0.22 | 0.23 | 0.08 | 0.04-0.42 | 0.87 | 0.88 | 0.07 | 0.69-1.29 |
| Males | All | 0.17 | 0.19 | 0.11 | 0.03-0.60 | 0.15 | 0.16 | 0.08 | -0.09-0.43 | 0.92 | 0.92 | 0.10 | 0.58-1.37 |
|  | Baseline | 0.14 | 0.19 | 0.13 | 0.03-0.54 | 0.16 | 0.16 | 0.10 | -0.09-0.43 | 0.90 | 0.93 | 0.12 | 0.65-1.37 |
|  | Day | 0.09 | 0.10 | 0.06 | 0.03-0.49 | 0.18 | 0.19 | 0.08 | 0.05-0.43 | 1.00 | 1.01 | 0.12 | 0.70-1.37 |
|  | Night | 0.31 | 0.31 | 0.10 | 0.06-0.52 | 0.12 | 0.11 | 0.09 | -0.09-0.39 | 0.85 | 0.85 | 0.06 | 0.65-1.22 |
|  | Cold | 0.24 | 0.25 | 0.10 | 0.10-0.60 | 0.14 | 0.15 | 0.07 | -0.05-0.39 | 0.91 | 0.90 | 0.07 | 0.71-1.18 |
|  | Day | 0.19 | 0.21 | 0.06 | 0.10-0.57 | 0.11 | 0.11 | 0.04 | -0.03-0.24 | 0.91 | 0.90 | 0.06 | 0.72-1.18 |
|  | Night | 0.31 | 0.32 | 0.10 | 0.12-0.60 | 0.18 | 0.19 | 0.07 | -0.04-0.39 | 0.92 | 0.91 | 0.07 | 0.71-1.15 |
|  | Hot | 0.11 | 0.13 | 0.08 | 0.03-0.47 | 0.17 | 0.17 | 0.08 | -0.04-0.43 | 0.94 | 0.94 | 0.10 | 0.58-1.34 |
|  | Day | 0.08 | 0.09 | 0.04 | 0.03-0.27 | 0.13 | 0.13 | 0.05 | -0.04-0.29 | 0.94 | 0.95 | 0.10 | 0.66-1.34 |
|  | Night | 0.18 | 0.19 | 0.10 | 0.04-0.47 | 0.23 | 0.23 | 0.07 | 0.04-0.43 | 0.93 | 0.93 | 0.09 | 0.58-1.30 |
|  | Data | VCO2 |  |  |  | VO2 |  |  |  |  |  |  |  |
|  |  | med. | mean | sd | range | med. | mean | sd | range |  |  |  |  |
| Females | All | 0.45 | 0.52 | 0.29 | 0.06-1.59 | 0.51 | 0.58 | 0.33 | 0.05-1.82 |  |  |  |  |
|  | Baseline | 0.38 | 0.46 | 0.28 | 0.07-1.59 | 0.41 | 0.51 | 0.34 | 0.05-1.59 |  |  |  |  |
|  | Day | 0.26 | 0.29 | 0.15 | 0.09-1.12 | 0.24 | 0.28 | 0.16 | 0.08-1.05 |  |  |  |  |
|  | Night | 0.64 | 0.69 | 0.26 | 0.10-1.39 | 0.78 | 0.81 | 0.30 | 0.14-1.59 |  |  |  |  |
|  | Cold | 0.65 | 0.69 | 0.28 | 0.17-1.59 | 0.71 | 0.76 | 0.30 | 0.19-1.82 |  |  |  |  |
|  | Day | 0.55 | 0.58 | 0.20 | 0.17-1.51 | 0.62 | 0.64 | 0.21 | 0.19-1.59 |  |  |  |  |
|  | Night | 0.82 | 0.87 | 0.30 | 0.17-1.59 | 0.89 | 0.95 | 0.33 | 0.19-1.82 |  |  |  |  |
|  | Hot | 0.33 | 0.39 | 0.22 | 0.06-1.24 | 0.38 | 0.45 | 0.25 | 0.06-1.44 |  |  |  |  |
|  | Day | 0.26 | 0.30 | 0.14 | 0.08-1.13 | 0.31 | 0.33 | 0.16 | 0.08-1.32 |  |  |  |  |
|  | Night | 0.49 | 0.53 | 0.24 | 0.06-1.24 | 0.57 | 0.61 | 0.28 | 0.06-1.38 |  |  |  |  |
| Males | All | 0.51 | 0.59 | 0.34 | 0.09-1.82 | 0.57 | 0.65 | 0.39 | 0.09-2.03 |  |  |  |  |
|  | Baseline | 0.43 | 0.58 | 0.36 | 0.10-1.62 | 0.48 | 0.66 | 0.43 | 0.09-1.85 |  |  |  |  |
|  | Day | 0.29 | 0.33 | 0.17 | 0.10-1.54 | 0.29 | 0.34 | 0.19 | 0.09-1.62 |  |  |  |  |
|  | Night | 0.90 | 0.89 | 0.30 | 0.10-1.57 | 1.07 | 1.04 | 0.33 | 0.20-1.79 |  |  |  |  |
|  | Cold | 0.72 | 0.76 | 0.29 | 0.34-1.82 | 0.80 | 0.85 | 0.32 | 0.33-2.03 |  |  |  |  |
|  | Day | 0.59 | 0.64 | 0.18 | 0.34-1.71 | 0.66 | 0.71 | 0.21 | 0.33-1.95 |  |  |  |  |
|  | Night | 0.95 | 0.97 | 0.30 | 0.36-1.82 | 1.05 | 1.08 | 0.34 | 0.39-2.03 |  |  |  |  |
|  | Hot | 0.34 | 0.41 | 0.26 | 0.09-1.42 | 0.36 | 0.45 | 0.28 | 0.10-1.58 |  |  |  |  |
|  | Day | 0.27 | 0.29 | 0.12 | 0.10-0.80 | 0.28 | 0.31 | 0.13 | 0.10-0.90 |  |  |  |  |
|  | Night | 0.56 | 0.60 | 0.30 | 0.13-1.42 | 0.61 | 0.65 | 0.32 | 0.12-1.58 |  |  |  |  |

Within each sex, the majority of pairwise comparisons of metabolic variables across treatment groups were significantly different (Fig. 2; Fig. S4). Separating each treatment into daytime and nighttime intervals found similar evidence of significantly different physiological responses among treatment groups, with few exceptions (Fig. 2; Fig. S5). Baseline daytime conditions were equivalent to the hot experimental conditions, yet metabolic responses unexpectedly differed between the two treatments for EE, RQ, and H_2_O (Table 2). As expected, male VO_2_ did not differ between cold and baseline nighttime treatments, during which the environmental conditions were identical, but these two time intervals did differed significantly in all other metabolic variables examined (Fig. S2). Both sexes exhibited their highest mean EE under constant cold conditions (Female = 0.23 J/s, Male = 0.25 J/s)and lowest under constant hot conditions (both = 0.13 J/s). Females lost less water under diurnally variable environmental conditions (mean = 0.12 mg), than they did under constant hot (0.19 mg) or cold (0.13 mg) treatments, while males lost less water under the constant cold treatment (0.15 mg), relative to baseline (0.16 mg) or hot (0.17 mg) treatments.

**Table 2.**
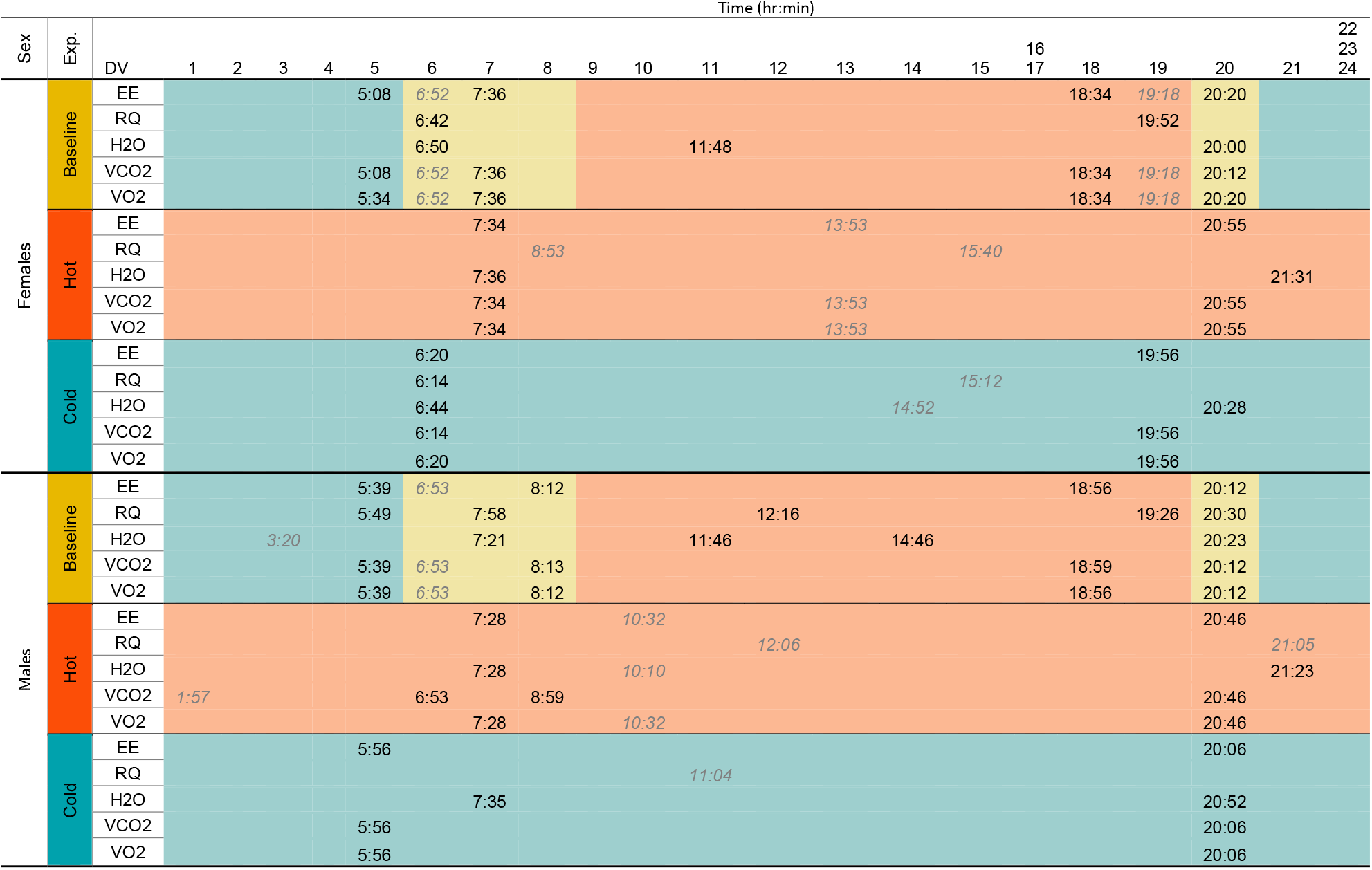
Maximum-likelihood change point estimates for males and females, across each experiment (baseline, hot, cold), based on AIC optimization. True change points in black, overfit change points in grey italics. Cold environmental conditions are denoted in blue, hot in orange, and environmental transitions (baseline only) are shown in yellow.

Weight was not correlated with any response variable (Fig. S3), therefore we did not include it as a covariate or run an ANCOVA. Our data violated assumptions of normality and homogeneity of variance required for an ANOVA (Tables S6, S7), so we ran nonparametric Kruskal-Wallis and Wilcox tests which identified significant differences among all but five pairs of group means (Fig. 2; Fig. S8). Female water loss did not differ significantly between the baseline and cold treatment groups; however, differences were detected when the data were split into nighttime and daytime intervals. Male EE did not differ between nighttime baseline and nighttime cold treatments (Fig. 2). The most water loss was observed under the hot treatment; however, peak water loss (0.43 mg for males, 0.37 mg for females) occurred under diurnally variable conditions. Mean EE was higher during the nighttime, relative to daytime, for all experiments.

### Signal processing of physiological time-series

Time-series data for EE, RQ, and H_2_O are visualized across a 24-hour cycle in Fig. 3 for all experiments and both sexes. Equivalent plots for VO_2_ and VCO_2_ are available in Fig. S9. Both sexes exhibited similar physiological patterning of EE across all treatment groups, but at different magnitudes of change (Fig. 3; Table 1). RQ was highly variable under diurnal environmental cycling, but flat-lined at the FQ (0.89) under both constant conditions (Fig. 3, 4). An RQ of one indicates exclusive use of carbohydrates, whereas an RQ of 0.8 indicates primarily protein consumption, and an RQ of 0.7 suggests the use of fats (Richardson, 1929). Under constant temperatures, RQ was fixed at 0.91 (+/− 0.07) for females and 0.90 (+/− 0.07) for males under hot conditions, and at 0.88 (+/− 0.07) and 0.94 (+/− 0.10) for females and males respectively, under constant cold conditions. Only under natural diurnally cycling, did RQ reach values greater than one, averaging 1.00 for males and 1.05 for females during the day and reaching a max of 1.37 and 1.50 for males and females, respectively. Patterns of water loss were inverted between baseline and constant treatments (Fig. 3). Under baseline conditions, water loss ramps up in the morning, decreases throughout the day and stabilizes at night. In contrast, under constant conditions, water loss mirrors EE patterning, decreasing during the day and increasing at night when animals resume higher levels of activity. Similar patterns are also observed in VO2 and VCO2 (Fig. 3).

**Figure 3.**
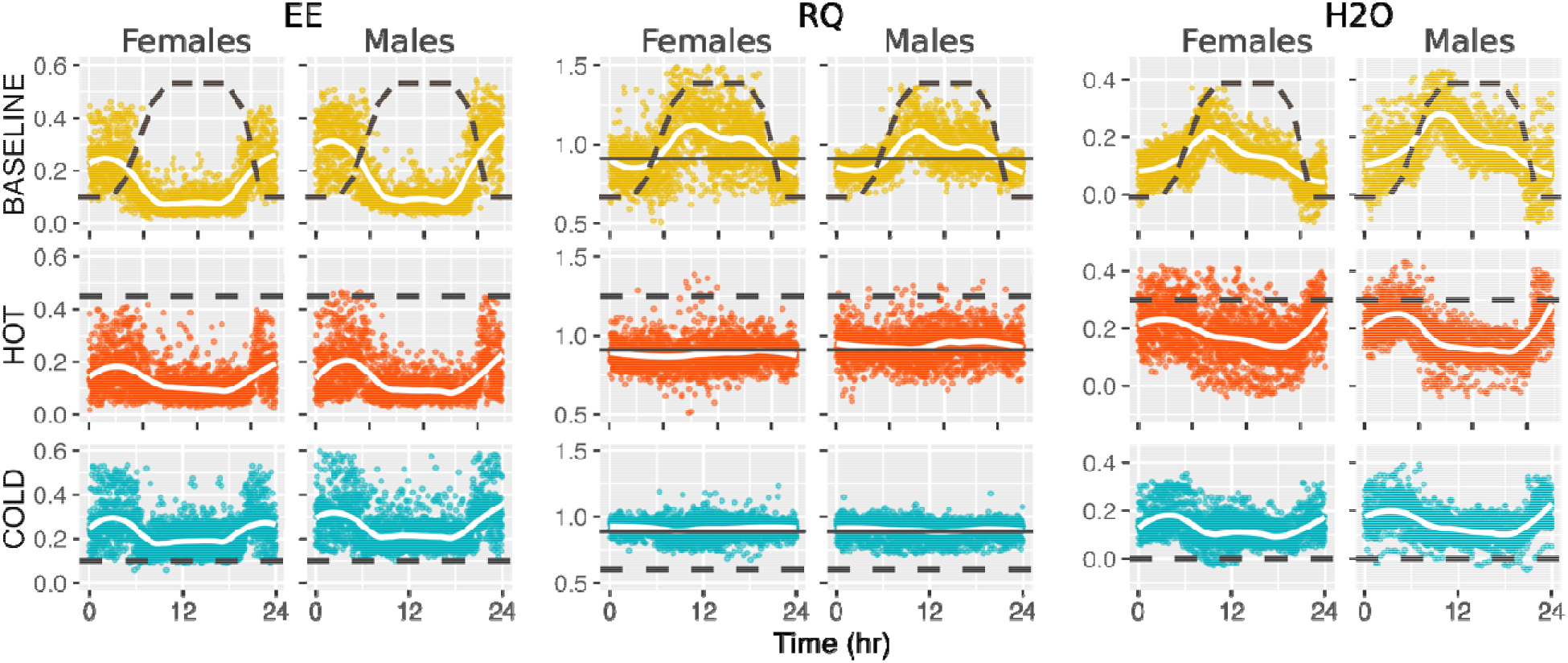
24-hour time series of energy expenditure (EE, left), respiratory quotient (RQ, center), and water loss (H_2_O, right) for males and females across the three experimental groups: yellow = baseline or normal diurnally variable environmental conditions, orange = constant hot, blue = constant cold. Black dashed lines depict scaled daily temperature variation across experiments and solid lines (RQ plots only) indicate the food quotient (0.89).

**Figure 4.**
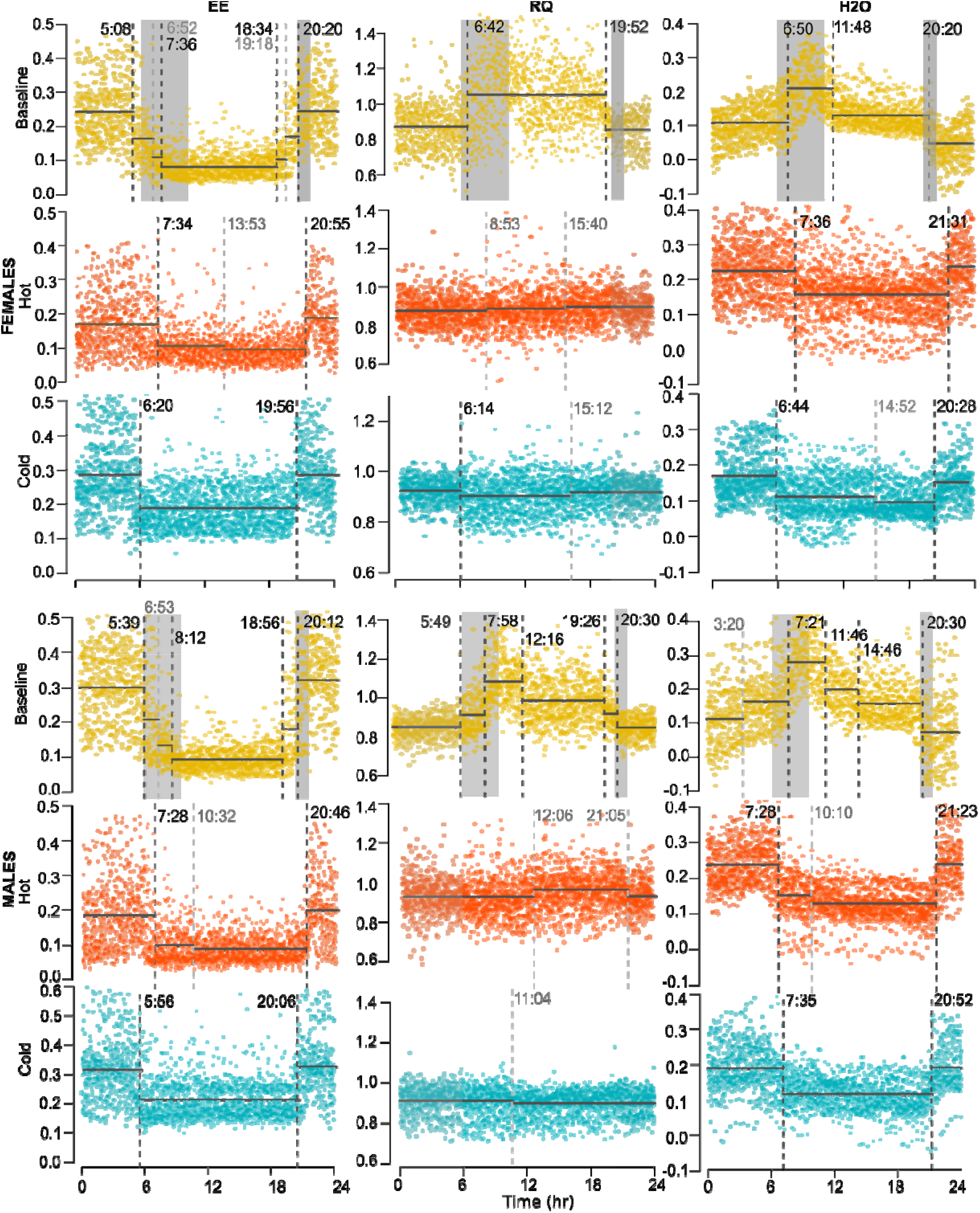
Change points (hr:min), denoting shifts in mean and variance for EE (left), RQ (center), and H_2_O (right) across a 24 hour cycle for females (top 3 rows) and males (bottom 3 rows) for each experiment (baseline = yellow, hot = orange, cold = blue). Vertical, dashed bars indicate change point estimates, with light grey indicating over fit change points. Light grey shading indicates nighttime (e.g., dark photoperiod), dark grey shading indicate transition periods (only present in baseline experimental treatments), and no colored shading indicates daytime (e.g., light photoperiod).

AIC optimized, maximum-likelihood change point estimates are in Table 2 and visualized in Fig. 4 (results for VO_2_ and VCO_2_ are visualized in Fig. S10). Under circadian environmental cycling, males exhibited an average of five change points and females three. As expected for change point analyses (e.g., fitting continuous data into a stepwise model), both sexes had change points overfit within rapid environmental transition periods (Fig. 4). Detected metabolic shifts roughly correspond to the initiation and conclusion of physical environmental transitions (morning, evening). Some additional change points were detected outside of environmental transition periods, including a large midday downshift in water loss present in both sexes. Females exhibit a morning shift in EE, VCO_2_, and VO_2_ earlier (05:16, on a 24-hour clock HH:MM) than males (05:41) on average. Morning change points for both sexes occurred prior to physical changes in temperature and humidity within the environment. Females experienced a second metabolic shift around 07:00 and males, just after 08:00. Evening metabolic shifts also occurred in two stages. Females experience an uptick in EE, VCO_2_, and O_2_ just after 18:30, and males just before 19:00, with a secondary shift around 20:12 for both sexes. We observed a single, large upward morning shift in water loss during the morning environmental transition. That initial uptick is followed by at least one midday downshift in water loss, followed by an additional downshift coincident with (20:00 females) or soon after (20:23 males) the initiation of the evening transition. Females exhibited only two change points in RQ, one during the morning transition, after the change in photoperiod and environment, and another preemptive of the evening transition. Males exhibited five AIC-optimal change points in RQ: one just before and one during each transition (morning, evening) and another midday coincident with the midday downshift in water loss. Means and standard deviations for each segment (i.e., period of mean and variance stability between estimated change points) are reported in Table S11 for both the optimal number of change points and excluding potentially overfit change points. Bayesian change point results (Table S12) are generally concordant with maximum-likelihood estimates.

Under constant hot conditions, both sexes experienced delayed (>2 hr on average) metabolic shifts relative to baseline and all major shifts occurred after changes in photoperiod. Under constant cold conditions, however, the morning transition occurred later than it had under baseline conditions for both sexes, but the shift remained preemptive of photoperiod change in males. Evening metabolic transitions under cold conditions also moved forward in time, but under this condition, the female shift occurred prior a change in photoperiod, while the male shift occurred after.

## DISCUSSION

In this study, we compared the metabolic responses of male and female cactus mice under diurnally variable and constant (hot and dry, cold and wet) conditions to assess the influence of temperature, humidity, and photoperiod on adaptive metabolic regulation. Overall, metabolic patterning was significantly shaped by environmental conditions, females were less heat tolerant than males, and metabolic responses were best optimized under diurnally variable conditions. An RQ greater than one, suggestive of lipogenesis, is observed but only under circadian environmental conditions, supporting the hypothesis that metabolism is well-tuned to environmental cycling. A number of these observations would not have been observed under constant environmental conditions and highlight the importance of continuous metabolic phenotyping through environmental transitions to better understand the physiological responses of organisms to dynamic environments.

### Metabolic sexual dimorphism

Although previous studies have found different results for sexual dimorphism in this species (females slightly outweigh males, Davis, 1966; no dimorphism, Manning et al., 2006), we did not detect body weight dimorphism between the sexes, nor a correlation between weight and any measured metabolic variable. Despite the lack of dimorphism, metabolic responses varied significantly between non-reproductive adult males and females (Figs. S1, S2). While less expected in the absence of size differences, metabolic demands frequently vary between sexes (Mittendorfer, 2005; McPherson and Chenoweth, 2012; Carmona-Alcocer et al., 2012) and may be explained by differences in behavior or activity level, for which EE represents a coarse proxy. In general, females spent less energy than males (Table 1; Fig. 2), resulting in corresponding differences in VO_2_, VCO_2_, and H_2_O, among other variables. Nonetheless, the overall patterns of variation, including the direction, amplitude, and frequency of change across metabolic response variables were similar among the sexes for all treatment groups (Fig. 3, 4).

Under normal diurnal cycling, respiratory water loss varied most between the sexes, with males losing significantly more water than females. This observation may be in part explained by higher activity levels in males (observed by not quantitatively reported herein), as increased activity levels lead to greater water loss through increased respiration (Bruck, 1962). Females used less energy than males during the daytime under environmentally variable conditions; Yet, under constant hot conditions, females lost more daytime water than males, despite similar levels of EE and identical environmental conditions across daytime diurnal and hot treatments (Table 1). This suggests that females may be less tolerant of extended heat stress. The opposite pattern is observed for males which appear to be less cold tolerant. Males spend significantly more energy relative to females under constant cold conditions (Table 1), likely as a consequence of increased thermogenesis. Consistent with this hypothesis, RQ is also lower for females under cold conditions, but higher under constant hot conditions (Table 1). Even though only non-reproductive individuals were examined in this study, differences in reproductive biology or sex-specific selection may have influenced adaptive evolution in this species. Sexual dimorphism in heat tolerance is consistent with observations in other mammals, including humans (Glucksman, 1974; Mittendorfer, 2005; Pomatto et al., 2018). While endothermy provides the advantages of a stable internal body temperature, which may differentially impact the fitness of males and females. For example, in males, a body temperature increase of only a few degrees centigrade above normothermia can reduce the viability of stored spermatozoa (Moore, 1926). This challenge is typically resolved through externalizing the testes to the scrotum, to maintain a lower local temperature. Although the non-reproductive cactus mice examined here hold their testes inside their abdominal cavity, like most other non-reproductive rodents. Even in cases of extreme heat, hyperthermia usually results in reduced fertility, albeit only temporarily, as spermatozoa are quickly replaced. Yet, the fitness consequences of hyperthermia may be greater in females, where the embryo cannot be thermally isolated and damage is irreversible (Greenwood and Wheeler, 1985). In consequence, the actions of sex-specific selection may lead to differences in traits that may not, initially, appear to be directly associated with sexual competitiveness, such as growth rate, thermoregulation, metabolic biorhythms, and environmental sensory (Glucksman, 1974; McPherson and Chenoweth, 2012; Juárez-Tapia and Miranda-Anaya, 2017; Calisi et al., 2018). Difference in lipid kinetics are hypothesized to maintain sexually dimorphic metabolic phenotypes (Mittendorfer, 2005). Similarly genes involved in lipid metabolism have been identified as important for thermoregulation in high-altitude adapted *Peromyscus* (Cheviron et al., 2012) and under selection in wild cactus mice (Tigano et al., 2020), reinforcing an important role for sex-specific lipid metabolism in the physiological adaptation of this species to desert environments.

### Circadian metabolic tuning

We hypothesized that desert-adapted cactus mice would exhibit metabolic responses that minimize water loss and limit EE under hot and dry environmental conditions. Indeed, we found significant support for environmentally-associated adaptive metabolic responses for all examined variables. Circadian patterning of EE is observed across all experimental treatments and is consistent with the nocturnal activity pattern of this species (Warner et al., 2010; Refinetti, 2010): Higher at night and lower during the day. Activity patterns, like nocturnality, are known to shape patterns of circadian metabolism and in mammals, shifts in activity are often governed by photoperiod (Miranda-Anaya et al. 2019). In deserts, increased locomotion and food intake at night minimizes evaporative water loss during hot and dry daytime conditions when physiological cooling is most challenging (Abreu-Vieira et al., 2015). Animals and, more specifically, near relatives of *Peromyscus* also maintain a warmer body temperature during the active (nighttime) phase relative to the inactive (daytime) phase that cannot be wholly explained by differences in activity levels, but may require additional metabolic resources to achieve (Abreu-Vieira et al., 2015). EE is the sum of an organism’s BMR, physical activity, and energy invested in cold-induced thermogenesis; therefore, EE represents only a coarse approximation for differences in activity level. At thermoneutrality, mice (*Mus*) expend approximately 60% of their total EE on BMR, 12% on the thermic effects of food, 25% on physical activity, and 0% on cold-induced thermogenesis (Abreu-Vieira et al., 2015). Thermogenesis (Storz et al., 2019; Bastias-Perez et al., 2020) likely explains both the elevated EE observed under constant cold conditions and the greater variance in EE observed under diurnally variable environmental conditions, where thermogenesis is expected to occur nightly but not during the day (Fig. 2; Table 1). Similar patterns of EE across all treatments (high at night and lower during the day) is consistent with the nocturnal activity pattern of this species and may be modulated in part by photoperiod, which remained constant across all treatment groups (Miranda-Anaya et al. 2019). The role of photoperiod in structuring activity patterns, and by proxy EE, remains to be experimentally explored for this species.

In a captive environment where food is not a limiting resource, animals spent less energy at night under constant hot conditions, than they did under diurnally variable conditions, suggesting that extreme or extended temperature events may shape foraging strategies or activity patterns into the future (Du Plessis et al., 2012; Mason et al., 2017; Riddell et al., 2019). Behavioral thermoregulation can be achieved through increasing or decreasing activity or by relocating to a thermally suitable microclimate (e.g., underground burrow). While behavioral thermoregulation is energetically less expensive than autonomic thermoregulation (Terrien et al., 2011; Hayford et al., 2015), there is a tradeoff between the energy and time invested in thermoregulation and that which remains available for other biological processes critical to survival. Increased investment in thermoregulation diverts time and energy away from reproduction and resource acquisition and physical relocation to a suitable microclimate costs energy and may drive animals further away from areas that are favorable in terms of resource availability or predation risk (Dunbar, 1998; Kearney and Porter, 2004; Mason et al., 2017).

Under diurnally variable environmental conditions, both sexes experienced greater variance in all metabolic response variables across a 24-hr period than under constant conditions, indicating that metabolic regulation is both dynamic and environmentally mediated. Cactus mice better optimize energy expenditure, water loss, and fuel resources under diurnally variable environmental conditions, relative to artificially constant conditions. For example, nighttime water loss was reduced under diurnal conditions, despite high activity levels, whereas the reverse was true for both constant temperature treatments: high nighttime water loss and EE. As expected, water loss was greatest under constant hot and dry conditions and lowest under constant cold and wet conditions (Table 1). However, when split into daytime and nighttime intervals, animals in the diurnal treatment group lost significantly less water (Fig. 1), even when conditions were identical (e.g., daytime diurnal and hot conditions). In fact, during the daytime under diurnally variable conditions, animals lost even less water than they did under constant cold conditions, suggesting the water economy of cactus mice is most efficient under natural diurnal cycling or when the animals are not thermally stressed. Although significant, the volumetric difference in water loss between treatments is small (0.01-0.07), yet the biological consequences of these differences remain to be tested in detail.

### Lipogenesis as potential a buffer against dehydration

The impact of environmental variation on cactus mouse metabolism is most obvious for RQ. Under diurnally variable conditions, RQ increased with increasing temperature and humidity, followed by an evening downshift, coincident with or occurring just prior to the evening transition to cooler, wetter conditions (Figs. 3, 4). In contrast, RQ flat-lined at the FQ under both constant conditions. This indicates that RQ in cactus mice is tightly modulated by temperature, and more specifically, sustained temperatures.

An RQ over one can indicate either activity levels above aerobic threshold (Whipp, 2007; Zagatto et al., 2012) or lipogenesis (Benedict and Lee, 1937; Kleiber, 1961; Elia and Livesey, 1988; Abreu-Vieira et al., 2015; Levin et al., 2017). For cactus mice, elevated RQ levels are observed during the daytime, when mice are inactive and are therefore suggestive of facultative lipogenesis. Glycolysis converts carbohydrates (glucose) into pyruvate, which is then converted into acetyl-CoA. Excess acetyl-CoA can be used to produce fatty acids and triglycerides through the process of lipogenesis. During this process, glycerol hydroxyl groups (-OH) react with the carboxyl end (-COOH) of a fatty acid chain to produce water (H_2_O) and carbon (C) atoms that then bind to oxygen (O) through a dehydration synthesis reaction, forming water and carbon dioxide (Ameer et al., 2014; OpenStax, 2016). In consequence, lipogenesis results in a measurable uptick in released CO2, as evidenced by a spike in RQ (the ratio of VCO2 to VO2). Since free water is scarce in desert environments, we hypothesize that the breakdown of food via lipogenesis may be important indirect source of water for cactus mice (Bradley and Yousef, 1972), whereby the endogenous synthesis of fatty acids and triglycerides may buffer against the negative physiological effects of dehydration. Interestingly, an RQ greater than one, consistent with lipogenesis, is only observed during the daytime under diurnally variable environmental conditions and surprisingly, never recorded under constant hot and dry conditions. Circadian cycling of lipogenesis is observed in other rodents (hamsters, rats, mice), modulated by dietary and hormonal variation (insulin and prolactin, Kimura et al., 1970; Cincotta and Meier, 1985; Bryson et al., 1993; Kersten, 2001; Letexier et al., 2003; Abreu-Vieira et al., 2015). Insulin-associated genes have been shown to be under selection in cactus mice *(Insl3, Igfbp3;* Kordonowy et al., 2017) and *Inpp5k* appears to be a functionally important transcript for water regulation in this species, that is also involved in insulin signaling (MacManes, 2017; Bridges and Saltiel, 2015).

Circadian patterning of RQ, and therefore also VCO_2_ and VO_2_, in cactus mice under environmentally variable conditions is likely what was described as ‘circadian torpor’ by Macmillen (1965, 1972). While the abiotic regulatory mechanism of lipogenesis in cactus mice remains elusive, it is not driven by photoperiod, nor constant temperature or humidity. Circadian lipogenesis also does not appear to be resource-dependent, as originally hypothesized by Macmillen (1965, 1972; also see Stephan, 2001), because animals were provided food and water *ad libitum* throughout the experiments. Potentially consistent with Macmillen’s (1965, 1972) resource-limitation hypothesis, however, RQ patterning suggests that lipogenesis may be glucose-limited, at least within a single day. The lipogenesis reaction is controlled by hepatic insulin levels (Ameer et al., 2014), thus, we hypothesize that lipogenesis decreases from morning to evening in response to decreasing insulin levels or leptin-ghrelin signaling (Buijs et al., 2017; Luna-Moreno et al., 2017), which are not mutually exclusive. For example, leptin regulates hepatic insulin sensitivity and glucose homeostasis, as well as energy expenditure (Inoue et al., 2004). Insulin levels are expected to become depleted over the course of the day, while nighttime refeeding is anticipated to increase serum insulin, and in turn, potentially stimulate lipogenesis (Kimura et al., 1970; Bryson et al., 1993; Yamajuku et al., 2012; Liu et al., 2013; Adamovich et al., 2014). The absence of an RQ greater than one under either constant treatment suggests that this pattern is not solely mediated by insulin availability. Extended exposure to sub-optimal environmental conditions can negatively impact body condition and survival in endotherms (McKechnie and Wolf, 2010; Gardner et al., 2016); therefore, we hypothesize that the detection of extended sub-optimal conditions triggers a downshift in metabolic resource consumption in response to physiological stress (Taborsky et al., 2020). In addition to the physiological limits of an organism, environmental stability is a key factor in shaping stress responses (Taborsky et al., 2020). Continuous metabolic phenotyping through state transitions (multiple days of diurnal cycling to multiple days of constant conditions) will be necessary to determine the abiotic triggers and metabolic processes responsible for circadian variation in RQ.

High altitude adapted *Peromyscus* exhibit metabolic differences that enhance survival in cold, hypoxic (Ivy and Scott, 2018, 2020) environments. In these species, cold-induced thermogenesis is used to maintain a constant internal body temperature and is regulated by flexibility in lipid metabolism (Cheviron et al., 2012; Munshi-South and Richardson, 2017). Lipids are also known to regulate circadian rhythmicity in mammals, with hepatic lipid metabolism being controlled by a suite of circadian Clock genes (Eckel-Mahan et al., 2012; Adamovich et al., 2014, 2015) some of which have been identified as under selection in cactus mice *(BMAL1, BMAL2;* Colella et al., *in press).* Under acute dehydration, cactus mice lose > 20% of their body weight without neurological, behavioral, or cardiovascular impairment, which would normally impact other mammals (Kordonowy et al., 2017). Developmental exposure to thermal extremes can also shape adaptive responses (Ivy and Scott, 2020) and the role of development in metabolic regulation of dehydration tolerance remains to be explored for this species.

### Timing of metabolic shifts

Analyses of continuous metabolic phenotypes confirmed that cactus mice physiology is well-tuned to circadian patterns of environmental variation. We expected there to be a physiological lag between the sensation of environmental change and a measurable shift in the circadian system (Aguilar-Roblero, 2015). For example, in related a Cricetid species, *Neotomodon alstoni,* increased activity levels are observed immediately following the initiation of the dark phase *(e.g.,* lights out; Miranda-Anaya et al. 2019). In contrast, under normal circadian cycling, most metabolic shifts in cactus mice occurred in advance of physical environmental change, or the perception thereof (Fig. 3, 4). Evidence of anticipatory metabolic shifts support the hypothesis that entrainment of biorhythms is adaptive in highly variable environments.

The timing of most metabolic shifts is disrupted under constant temperature conditions; thus, circadian metabolic patterning is not exclusively regulated by access to food, temperature, or humidity. The persistence of two major metabolic shifts under both constant treatments, however, suggests that photoperiod may be at least in part responsible for maintaining biorhythms. Photic synchronization is common among mammals, in which the hypothalamus acts as a circadian pacemaker, reactive to ambient light (Miranda-Anaya et al. 2019). Observed temporal changes in metabolic shifts between baseline and constant treatment groups were consistent with males being more heat tolerant than females. Relative to males, females use less energy and lose less water under cold conditions, but experienced a longer period of nighttime activity under hot conditions. Tests of inverted day and night conditions will be necessary to test the strength of the observed circadian rhythm and role of photoperiod.

### Conclusions

In conclusion, our results highlight an important role for circadian environmental variation in the evolution of metabolic efficiency in desert-adapted cactus mice. Lipid metabolism is increasingly emerging as a common mechanism of circadian and thermoregulatory plasticity, that warrants further experimental investigation in this species through both dietary and environmental manipulation and *in vitro* tests of metabolic fuel usage (Adamovich et al., 2014). Studies of physiological in nontraditional model organisms (Bedford and Hoekstra 2015) provides a comparative framework for identifying shared physiological responses and mechanisms among species. Further, linking molecular mechanisms to the physiological phenomena described here will allow us to better characterize the role of genetics, environment, and behavior on complex physiological phenotypes (Partch et al., 2014).

## ACKNOWLEDGEMENTS

We thank all members of the MacManes Laboratory for useful comments on early versions of the manuscript; the Animal Resources Office and veterinary care staff at the University of New Hampshire; A. Gerson at the University of Massachusetts Amherst for analytic guidance and technical support; Z. Cheviron at the University of Montana; B. Joos and J. Klok at Sable Systems International for their guidance in setting up our respirometry system; and A. Westbrook for computational support. This work was supported by the National Institute of Health National Institute of General Medical Sciences [1R35GM128843 to M.D.M.].

## COMPETING INTERESTS

No competing interests declared.

## DATA AVAILABILITY

Raw ExpeData (SSI) files are available through DataDryad: https://doi.org/10.5061/dryad.f4qrfj6v0. Macro processing files, respirometry data, and cage sampling scheme files are available also on Data Dryad. All R scripts used in this project are available through GitHub at: github.com/jpcolella/peer_respo.

